# Old players in new posts: the role of P53, ATM and DNAPK in DNA damage-related ubiquitylation-dependent removal of S2P RNAPII

**DOI:** 10.1101/2021.02.22.432201

**Authors:** Barbara N Borsos, Vasiliki Pantazi, Zoltán G Páhi, Hajnalka Majoros, Zsuzsanna Ujfaludi, Ivett Berzsenyi, Tibor Pankotai

**Affiliations:** Department of Pathology, Faculty of Medicine, University of Szeged, 1 Állomás Street H-6725 Szeged, Hungary; (B.N.B.); (V.P.); (Z.G.P.); (H.M.); (Z.U.); (I.B.)

**Keywords:** P53, ATM, DNAPK, ubiquitylation, S2P RNAPII, E3 ligases, WWP2, CUL3, transcription block, DNA repair

## Abstract

DNA double-strand breaks are the most deleterious lesions for the cells, therefore understanding the macromolecular interactions in the DNA repair-related mechanisms is essential. DNA damage triggers transcription silencing at the damage site, leading to the removal of the elongating RNA polymerase II (S2P RNAPII) from this locus, which provides accessibility for the repair factors to the lesion. Ataxia-telangiectasia mutated (ATM) and DNA-dependent protein kinase (DNAPK) are the two main regulatory kinases of homologous recombination and non-homologous end joining, respectively. Although these kinases are involved in the activation of different repair pathways, they have common target proteins, such as P53. We previously demonstrated that following transcription block, P53 plays a pivotal role in transcription elongation process by interacting with S2P RNAPII. In the current study, we reveal that P53, ATM and DNAPK are involved in the fine-tune regulation of the ubiquitin-proteasome system-related degradation of S2P RNAPII. However, they act differently in this process: P53 delays the removal of S2P RNAPII, while ATM and DNAPK participate in the activation of members of E3 ligase complexes involved in the ubiquitylation of S2P RNAPII. We also demonstrate that WW domain-containing protein 2 (WWP2) and Cullin-3 (CUL3) are interaction partners of S2P RNAPII, thus forming a complex with the transcribing RNAPII complex.

**Simple Summary:** To ensure the proper repair following DNA double-strand breaks, the eviction of the arrested elongating RNA polymerase II (S2P RNAPII) is required. Here, we report an emerging role of P53, Ataxia-telangiectasia mutated (ATM) and DNA-dependent protein kinase (DNAPK) in the ubiquitin-proteasome system-dependent removal of S2P RNAPII. We also identified interactions between S2P RNAPII and WW domain-containing protein 2 (WWP2) or Cullin-3 (CUL3) (members of E3 ligase complexes), which are involved in the ubiquitylation of S2P RNAPII following DNA damage. Furthermore, the RNAPII-E3 ligase complex interactions are mediated by P53, ATM and DNAPK, which suggests potential participation of all three proteins in the effective resolution of transcription block at the damage site. Altogether, our results provide a better comprehension of the molecular background of transcription elongation block-related DNA repair processes and highlight an indispensable function of P53, ATM and DNAPK in these mechanisms.

## 1. Introduction

DNA double-strand breaks (DSBs) are the most deleterious lesions, thus the fine-tuning of the related repair processes is indispensable to prevent genome instability. Ataxia-telangiectasia mutated (ATM) kinase and DNA-dependent protein kinase (DNAPK) are principal regulators in the precise coordination of the two main subpathways of DSB repair, homologous recombination (HR) and non-homologous end joining (NHEJ), respectively [1–4]. Although ATM and DNAPK are responsible for the activation of different repair pathways, they have common target proteins, such as H2A.X and P53 [5–9]. Following DNA damage, ATM and DNAPK can phosphorylate P53 at Ser15, resulting in its activation and nuclear accumulation [6–9]. Recently it has been shown that upon ionizing radiation (IR), the initial protein level of P53 and S1981P (phosphorylation at Ser1981) ATM was dramatically increased following treatment with DNAPK inhibitor, which is the consequence of the suppression of DNAPK-mediated inhibitory phosphorylation on several residues of ATM [10–12]. These findings presume that following IR, ATM is the main activator kinase of P53. Intriguingly, as a response to IR, ATM can contribute to the activation of DNAPK via phosphorylation at Thr2609, assuming the stimulating attendance of ATM in NHEJ [13].

P53 is a well-known tumor suppressor, which function during transcription was used to be restricted to the initiation phase. However, a novel function of P53 in transcription elongation has been recently revealed in yeast and human models [14–18]. We demonstrated for the first time in human cells that P53 bound to non-sequence specific gene regions, which was further enhanced following Actinomycin D (ActD)-induced transcription block. Nonetheless, a significant decrease in RNA polymerase II (RNAPII) occupancy was detected upon ActD and interaction was established between P53 and the elongating RNAPII (S2P RNAPII), suggesting a possible role of P53 in the damage-related removal of RNAPII. The remission in S2P RNAPII level was proved to be the consequence of its proteasomal-mediated degradation. Moreover, P53 was found to co-localize with γH2A.X at the damage foci, which suggested a potential role of P53 in transcription silencing to allow the recruitment of DNA repair factors to the damage site [17].

According to the severity of the DNA damage, the fate of the stalled RNAPII can vary [19–21]. Cockayne syndrome group B (CSB) and Transcription factor II H (TFIIH) participate in the forward (at minor lesions) and reverse (at bulky lesions) translocation of the RNAPII, respectively [22,23]. However, upon serious damage, the permanently stalled S2P RNAPII is marked for polyubiquitylation-mediated proteasomal degradation to promote the efficient repair mechanism [24,25]. Following DNA damage, RNAPII remains to be phosphorylated at Ser2 to prevent the initiation of a new transcription cycle and allows its ubiquitylation-related removal [26]. Several E3 ligase complexes are involved in the damage-related ubiquitylation of S2P RNAPII, such as Neural precursor cell expressed developmentally down-regulated protein 4 (NEDD4), Breast cancer 1 (BRCA1)-BRC A1-associated RING domain protein 1 (BARD1) and ElonginA/B/C-Cullin-5-RING-box protein 2 (EloA/B/C-CUL5-RBX2) [27–30]. WW domain-containing protein 2 (WWP2) has been identified hitherto as an interaction partner of RNAPII and as a key E3 ligase in the DSB-related polyubiquitylation of RNAPII [31,32]. Moreover, DNAPK has been shown to have a potential role in transcription silencing by facilitating the WWP2-dependent ubiquitylation of S2P RNAPII and the recruitment of the 26S proteasome to the break site [21,31,33].

Here, we shed light on a potential involvement of P53 in the polyubiquitylation and subsequent removal of the stalled S2P RNAPII following ActD-induced transcription block. Furthermore, we demonstrate interactions between members of E3 ligase complexes [WWP2 and Cullin-3 (CUL3)] and S2P RNAPII. Aside from P53, these interactions and the ubiquitylation of S2P RNAPII are also mediated by the kinase activity of ATM and DNAPK. P53 negatively affects the removal of S2P RNAPII, suggesting an auxiliary role of P53 in transcription elongation. As a response to ActD, at later time-points, we detected less ubiquitylated S2P RNAPII both in the absence of P53 and in the loss of activity of either ATM or DNAPK. Nonetheless, our assumption is that P53 can act differently from both kinases since we also demonstrated that P53 hinders the early ubiquitylation of S2P RNAPII following ActD. While following ActD-induced transcription arrest, the activity of ATM and DNAPK is necessary for the induction of WWP2 E3 ligase, P53 delays the cellular response related to the ubiquitylation-mediated proteasomal degradation of S2P RNAPII.

## 2. Results

### 2.1. P53 delays the removal of S2P RNAPII from actively transcribed coding regions as a response to transcription block

We previously demonstrated that human P53 associates to transcriptionally active gene regions by a non-sequence specific manner. We found opposite binding patterns of P53 and RNA polymerase II (RNAPII) as a response to Actinomycin D (ActD)-induced transcription elongation arrest: while P53 showed increased binding, a reduction was observed in RNAPII occupancy, suggesting its directional chromatin removal [17].

To study whether P53 plays a role in the chromatin dislodgement of the arrested elongating RNA polymerase II (S2P RNAPII) following DNA double-strand break (DSB) induction, we monitored the profile changes of the S2P RNAPII by chromatin immunoprecipitation (ChIP) experiment on HCT116 p53+/+ and HCT116 p53-/- colorectal cancer cell lines. For this, we treated both cell lines with the transcription elongation blocking agent ActD for 6 h and 24 h. The occupancy of S2P RNAPII was tracked at an intronic and two exonic regions of two actively transcribed genes, *ACTB* and *CDK12* (Figure 1A and B, respectively). As a negative control, we used primers for an intergenic region, where active transcription does not take place (Figure 1C).

**Figure 1.**
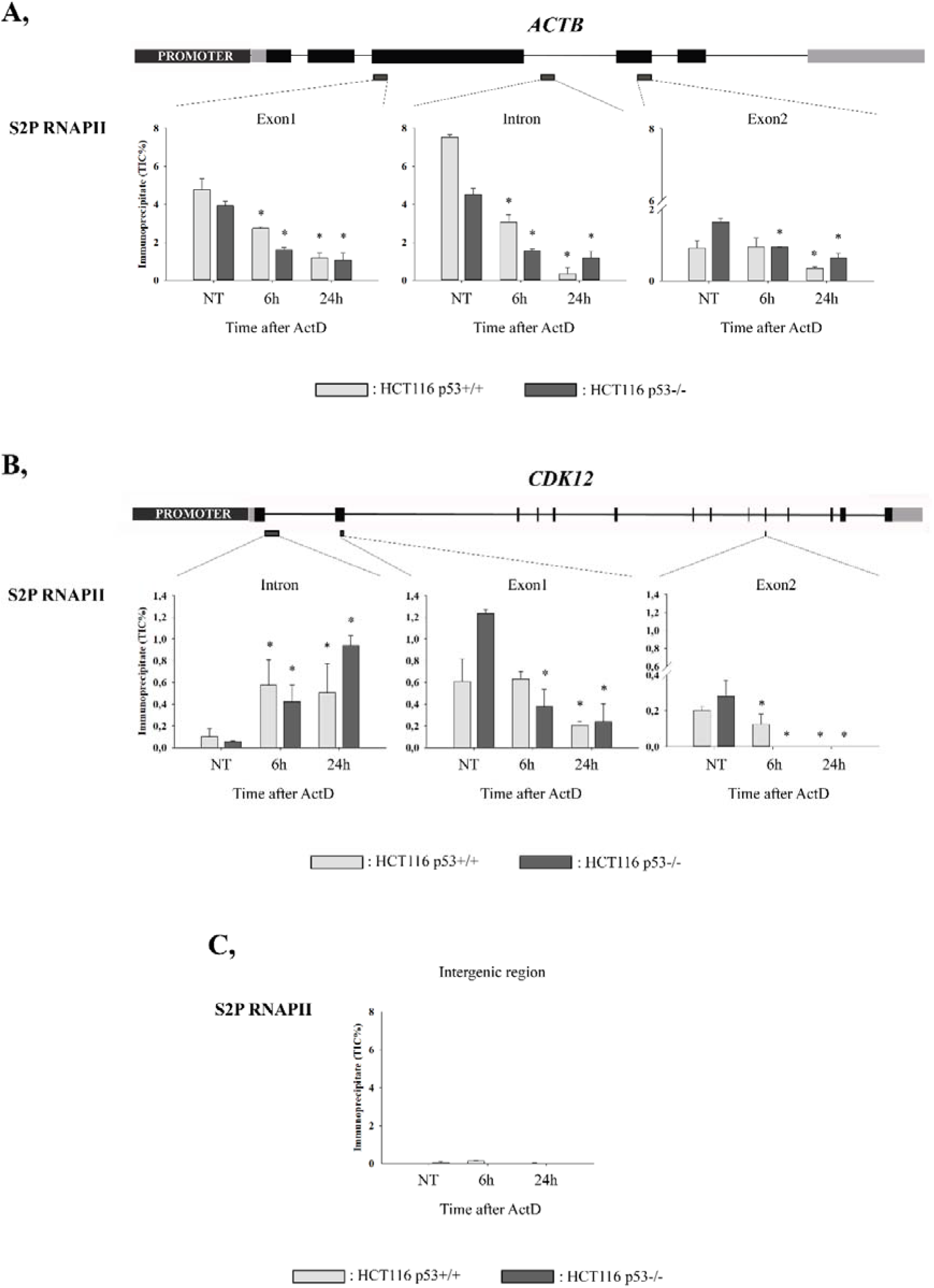
P53 affects the profile changes of elongating RNA polymerase II (S2P RNAPII) at transcriptionally active gene regions following Actinomycin D (ActD)-induced transcription elongation block. **(A-B)** S2P RNAPII occupancy was monitored with ChIP-qPCR at an intronic (Intron) and two exonic (Exon1 and Exon2) regions of *ACTB* and *CDK12* genes in the presence (light grey columns; HCT116 p53+/+ cell line) and in the absence (dark grey columns; HCT116 p53-/- cell line) of P53. The profile changes were tracked under physiological conditions (NT) as well as following 6 h and 24 h ActD treatment. (C) Primers designed to an intergenic region were used as the negative control of the ChIP. The schematic representation of the gene structures is indicated, and below the approximate chromosomal location of the qPCR amplicons is labelled with small, dark grey rectangles. The figure shows the representative result of one out of two independent experimental replicates. qPCR reactions were performed in triplicates. Stars represent statistical significance (*P ≤ 0.05) between the mean values. Mean values of ActD treated samples were compared to the mean value of the corresponding non-treated sample by one-way ANOVA in case of each cell line.

In HCT116 p53+/+ cells, S2P RNAPII occupancy is significantly reduced after 24 h ActD treatment, while in HCT116 p53-/- cells this attenuation can be observed already at 6 h. Nonetheless, in HCT116 p53+/+ cells, the binding of S2P RNAPII still remains high at 6 h, which denotes the stalled S2P RNAPII cannot be removed until this time-point in the presence of P53. Following 24 h ActD treatment, the remission in the occupancy of S2P RNAPII can be detected in both HCT116 p53+/+ and HCT116 p53-/- cell lines, suggesting a negative role of P53 in S2P RNAPII removal at earlier response during transcription elongation arrest. However, this phenomenon can be observed only at exonic regions. At the intronic region of either *ACTB* or *CDK12*, no remarkable differences can be detected in the binding tendency between the two cell lines (Figure 1A and B). These results reveal that in colorectal carcinoma cells, P53 delays the removal of the S2P RNAPII from actively transcribed exons as a response to transcription elongation block-induced DNA damage.

### 2.2. P53, ATM and DNAPK regulate the ubiquitylation of S2P RNAPII following ActD-induced transcription block

Following severe DNA damage, the elongating form of RNAPII is assigned to ubiquitylation-mediated proteasomal degradation to allow access for the recruitment of repair factors [21,24]. Using HCT116 p53+/+ and HCT116 p53-/- cell lysates, we pulled-down the ubiquitylated protein pool by tandem ubiquitin-binding entities (TUBEs). Subsequently, with immunoblot we examined the ubiquitylated S2P RNAPII (ub-S2P RNAPII) pool following 8 h and 24 h ActD treatment. To study the role of ATM and DNAPK in the ubiquitylation of S2P RNAPII following transcription elongation arrest, we applied ATM and DNAPK inhibitors (ATMi and DNAPKi), respectively. Ub-S2P RNAPII can be detected mainly at 8 h ActD treatment in each case, while at 24 h the majority of the ub-S2P RNAPII pool is presumably degraded by the 26S proteasome (Figure 2A and Figure S2). Intriguingly, in HCT116 p53-/- cells, much less ub-S2P RNAPII can be observed at 8 h ActD compared to that of detected in HCT116 p53+/+ cells (Figure 2A, lane 2 on right panel compared to lane 2 on left panel, respectively and Figure S2). Furthermore, regardless of the P53 status of the cell line, upon ATMi or DNAPKi treatment, also less ub-S2P RNAPII can be detected at 8 h compared to the corresponding sample in which no inhibitor was applied (Figure 2A, lane 5 and 8 compared to lane 2 on left and right panels and Figure S2).

**Figure 2.**
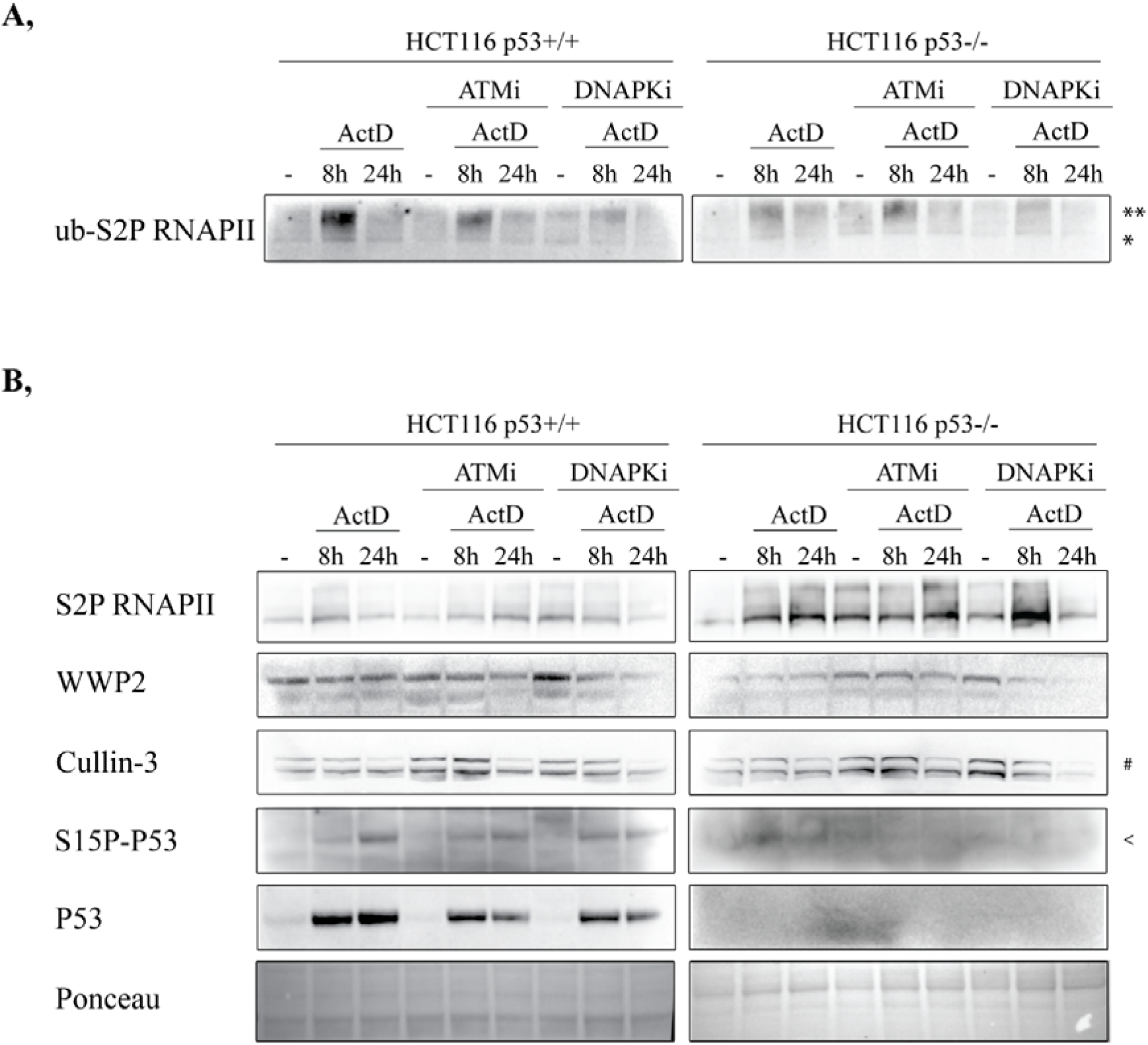
P53, Ataxia-telangiectasia mutated (ATM) and DNA-dependent protein kinase (DNAPK) influence the ubiquitylation of S2P RNAPII as a response to ActD treatment. (**A**) Tandem ubiquitin-binding entities (TUBEs) pull-down, followed by Western blot detection was performed in HCT116 p53+/+ (left panel) and p53-/- (right panel) cells. TUBEs experiment was accomplished under physiological conditions, 8 h and 24 h following ActD treatment in the presence or in the absence of ATMi or DNAPKi. From the pulled-down polyubiquitylated protein pool, polyubiquitylated S2P RNAPII (ub-S2P RNAPII) was detected with anti-S2P RNAPII antibody. *: monoubiquitylated S2P RNAPII, **: polyubiquitylated S2P RNAPII (**B**) Western blot experiment on whole cell lysates (referred to as input samples) of HCT116 p53+/+ (left panel) and p53-/- (right panel), which were used for TUBEs pull-down assay. Protein level changes of S2P RNAPII, WW domain-containing protein 2 (WWP2), Cullin-3, S15P-P53 and P53 following 8 h and 24 h ActD treatment in the presence or in the absence of ATMi or DNAPKi were immunodetected using specific antibodies. Ponceau staining was applied to detect the equal loading of the input samples. #: NEDD8-conjugated Cullin-3, <: unspecific bands of S15P-P53.

In HCT116 p53+/+ cells, the total protein level of S2P RNAPII is increased at 8 h ActD, then it returns to its normal level after 24 h (Figure 2B, first row, lane 1-3 on left panel and Figure S2). On the contrary, in HCT116 p53-/- cells the S2P RNAPII level still remains accumulated after 24 h ActD treatment (Figure 2B, first row, lane 1-3 on right panel and Figure S2). Strikingly, ATMi and DNAPKi treatments result in a reciprocal effect on S2P RNAPII protein level in both cell lines. In HCT116 p53+/+, S2P RNAPII is temporally accumulated following the combined treatment with ATMi and ActD, while it is getting reduced upon DNAPKi and ActD co-treatment (Figure 2B, first row, lane 4-6 and lane 7-9 on left panel, respectively and Figure S2). In HCT116 p53-/- cells, a slight decrease can be observed as a response to ATMi and 8 h ActD, while at the same time-point a dramatic elevation is detected in S2P RNAPII protein level following DNAPKi (Figure 2B, first row, lane 4-5 and lane 7-8 on right panel, respectively and Figure S2). From these data, it seems that following ActD-induced transcription block, the activity of either ATM or DNAPK influences the S2P RNAPII level in a P53-dependent manner.

Previously it was demonstrated in human cells that WW domain-containing protein 2 (WWP2) ubiquitylates S2P RNAPII, resulting in DNAPK-dependent transcription silencing and the subsequent activation of NHEJ pathway [31]. To reveal whether the WWP2 protein level is affected by ATM or DNAPK following transcription arrest, we applied ATMi or DNAPKi on ActD-treated HCT116 p53+/+ and HCT116 p53-/- cells, respectively (Figure 2B, second row on both left and right panels and Figure S2). As a result, upon DNAPKi treatment, WWP2 protein level is getting decreased following 8 h and 24 h ActD regardless of the presence of P53 (Figure 2B, second row, lane 7-9, on both left and right panels and Figure S2). ATMi can slightly reduce the WWP2 protein level after only 24 h ActD treatment in both cell lines (Figure 2B, second row, lane 6, on both left and right panels and Figure S2). However, WWP2 protein level is not affected by solely ActD treatment (in correlation with previously published results [31,32]), much less protein can be detected in the absence of P53 irrespectively of the cellular conditions (Figure 2B, second row, lane 1-3, on right panel compared to that of detected on the left panel and Figure S2). In HCT116 p53-/- cells, in basal conditions, this low WWP2 level is compensated both upon ATMi and DNAPKi (Figure 2B, second row, lane 1, 4, 7, on right panel and Figure S2). Furthermore, this phenomenon can be observed even at 8 h ActD treatment following ATMi (Figure 2B, second row, lane 5, on right panel and Figure S2). Apart from WWP2, we also monitored the changes in Cullin-3 (CUL3; a scaffold of another E3 ligase complex) protein level (Figure 2B, third row and Figure S2). CUL3 is part of the BCR (BTB-CUL3-RBX1) E3 ubiquitin ligase complex, which can become activated upon binding a neddylated (NEDD8-conjugated) CUL3 capable of heterodimerization [34]. We can detect the same pattern in CUL3 as in case of WWP2 following the above described conditions (Figure 2B, third row compared to second row, on both left and right panels and Figure S2). The neddylated and activated form of CUL3 is decreased following 24 h ActD treatment in both cell lines (Figure 2B, third row, upper lane 3,6,9, on both left and right panels and Figure S2). Although in contrast to WWP2, CUL3 protein level is not affected by the P53 status of the cells. Nonetheless, aside from DNAPK, ATM plays a minor role in the activation of WWP2 and CUL3, the main regulatory kinase of them is presumably the DNAPK.

Additionally, we investigated the changes in the protein level of P53 and its S15P form (S15P-P53) upon ActD, ATMi and DNAPKi (Figure 2B, fifth and fourth rows left and right panels, respectively and Figure S2). We detected an increase in the protein level of the total P53 pool and also its S15P form following transcription elongation block (Figure 2B, fifth and fourth rows, lane 1-3, on left panel, respectively and Figure S2). As expected, applying ATMi and DNAPKi led to a milder accumulation and activation of P53, supporting the efficiency of both inhibitors (Figure 2B, fifth and fourth rows, lane 5-6 and lane 8-9 on left panel, respectively and Figure S2).

These results suggest a pivotal role of P53, ATM and DNAPK in the ubiquitylation of S2P RNAPII following transcription elongation arrest. Nonetheless, based on the ChIP result represented in Figure 1 and the changes in WWP2 and CUL3 protein levels as a response to ActD treatment, P53 might be involved in different cellular processes than ATM and DNAPK. In the absence of P53, the chromatin removal of S2P RNAPII can be observed already at 6 h ActD (seen in Figure 1A-B), while at 8 h much less polyubiquitylated S2P RNAPII can be observed (Figure 2A and Figure S2), which suggests its pre-ubiquitylation at earlier time-points. In conclusion, following transcription elongation block, P53 presumably delays the ubiquitylation and subsequent proteasomal degradation of S2P RNAPII, while ATM and DNAPK might facilitate the activation of E3 ligases involved in the ubiquitylation of S2P RNAPII.

### 2.3. P53 counterarts the early ubiquitylation of S2P RNAPII following transcription elongation arrest

To further confirm our hypothesis that P53 is involved in the early ubiquitylation-mediated removal of S2P RNAPII, we first monitored the alterations in S2P RNAPII level following 1-8 h ActD treatment. In the presence of P53, we detected a constant temporal accrual in S2P RNAPII level. While in the absence of P53, the S2P RNAPII level is vagarious following ActD treatment: the initially accumulated S2P RNAPII was then reduced, while at 7 h and 8 h S2P RNAPII level was increased (Figure S1). Since in ChIP results shown in Figure 1A-B, we found that in the absence of P53, the occupancy of the stalled S2P RNAPII is highly reduced after 6 h and is further attenuated at 24 h, the detected accrual in the protein level of S2P RNAPII cannot be the stalled polymerase subpopulation, but rather those polymerases, which are accumulated at DNA ends to participate in *de novo* transcription and hence facilitate the proper repair process.

Based on this preliminary Western blot experiment, we selected three time-points (1 h, 4 h and 8 h) following ActD treatment at which we investigated the amount of ub-S2P RNAPII. For this, we performed TUBE assay in 1 h, 4 h and 8 h ActD treated as well as non-treated HCT116 p53+/+ and p53-/- cell lysates and by a subsequent immunoblot detection we compared the level of polyubiquitylated S2P RNAPII among these samples (Figure 3A and Figure S3). In HCT116 p53+/+ cells, we detected elevated level of polyubiquitylated S2P RNAPII following 8 h ActD treatment, while in p53-/- cells much less polyubiquitylated S2P RNAPII can be observed, supporting our previous results represented in Figure 2A (Figure 3A, lane 4 and 8, respectively and Figure S3). Intriguingly, at 1 h and 4 h post-ActD treatment only a limited amount of polyubiquitylated S2P RNAPII can be seen in the presence of P53 (Figure 3A, lane 2-3 and Figure S3). While in the absence of P53, as a response to 1 h ActD treatment, a relatively high amount of polyubiquitylated S2P RNAPII is detected, which supports our hypothesis that P53 plays a potential negative role in the early ubiquitylation of S2P RNAPII (Figure 3A, lane 6 and Figure S3). Therefore we believe that P53 postpones the dislodgement of S2P RNAPII from the damaged chromatin. Monoubiquitylated S2P RNAPII (lower lane) can be detected in each sample, including the basal conditions, as it is a crucial step even for the normal transcription cycle, which does not necessarily result in proteasomal degradation. Furthermore, monoubiquitylation is indispensable for the further polyubiquitylation of S2P RNAPII, thus its level is relatively high under stress conditions (as it is seen in Figure 3A).

**Figure 3.**
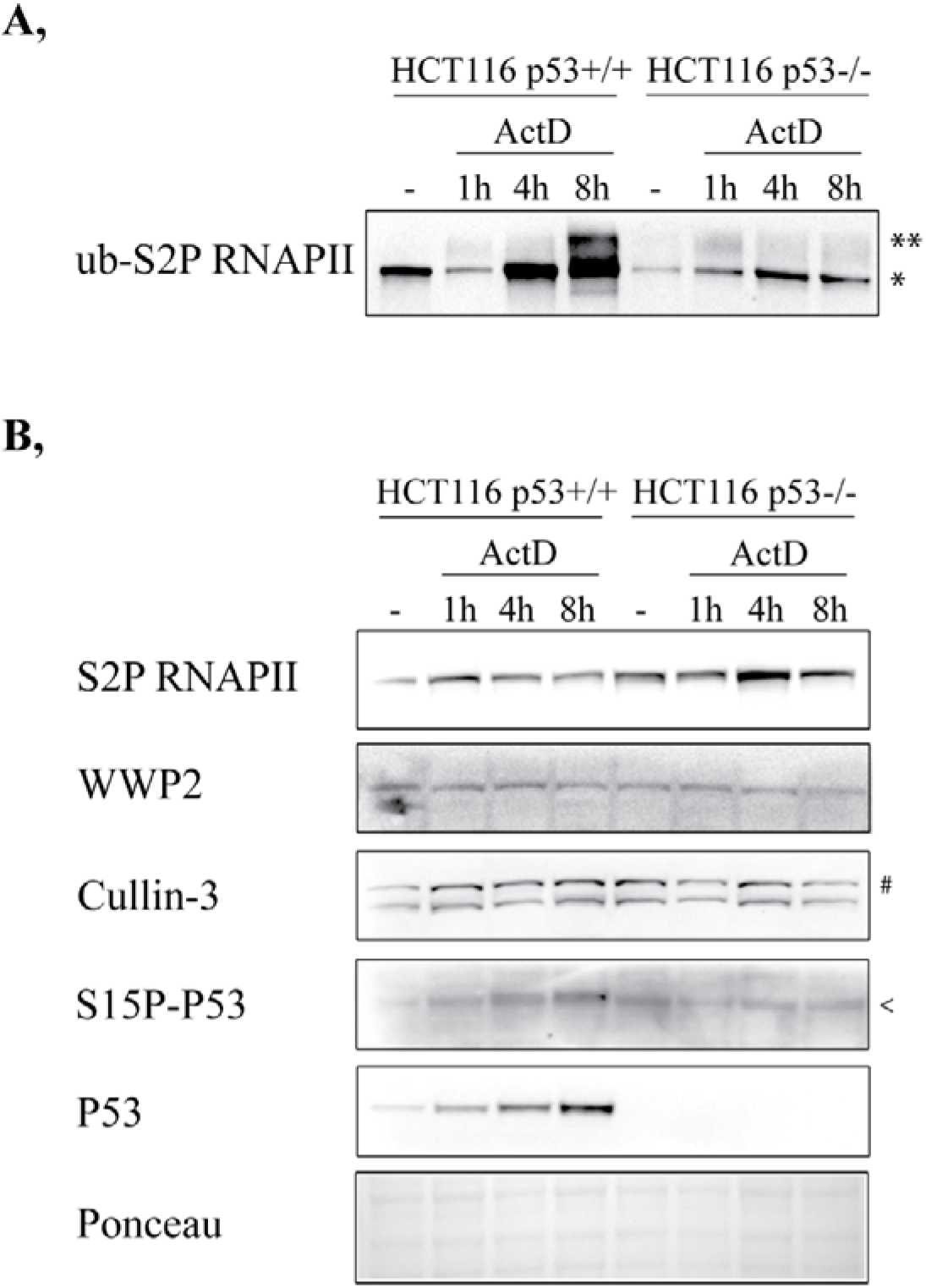
P53 is essential for the early ubiquitylation of RNAPII following ActD treatment. **(A)** Tandem ubiquitin-binding entities (TUBEs) pull-down, followed by Western blot detection was performed in HCT116 p53+/+ and p53-/- cells. TUBEs experiment was accomplished under basal conditions as well as 1 h, 4 h and 8 h following ActD. From the pulled-down polyubiquitylated protein pool, polyubiquitylated S2P RNAPII (ub-S2P RNAPII) was detected with anti-S2P RNAPII antibody. *: monoubiquitylated S2P RNAPII, **:polyubiquitylated S2P RNAPII. **(B)** Western blot experiment on whole cell lysates (referred to as input samples) of HCT116 p53+/+ and p53-/-, which were used for TUBEs pull-down assay. Protein level changes of S2P RNAPII, WW domain-containing protein 2 (WWP2), Cullin-3, S15P-P53 and P53 following 1 h, 4 h and 8 h ActD treatment were immunodetected using specific antibodies. Ponceau staining was applied to detect the equal loading of the input samples. #: NEDD8-conjugated Cullin-3, <: unspecific bands of S15P-P53.

Additionally, we monitored the alterations in the protein level of S2P RNAPII, WWP2, CUL3, S15P-P53 and P53 in the input samples of the TUBE pull-down (Figure 3B and Figure S3). In HCT116 p53+/+ cells, S2P RNAPII protein level was increased following 1 h, 4 h and 8 h ActD treatment (Figure 3B, first row, lane 1-4 and Figure S3). In HCT116 p53-/- cells, a significant accrual can be detected at 4 h (Figure 3B, first row, lane 7 and Figure S3). As the polyubiquitylated S2P RNAPII has been already degraded by that time-point, it cannot be the ub-S2P RNAPII pool, but it rather refers to an increased rate of *de novo* transcription during the repair process. WWP2 and CUL3 protein levels are not significantly changed at the examined time-points (Figure 3B, second and third row and Figure S3). In HCT116 p53+/+ cells, the level of P53 and its S15P form was getting increased in a time-dependent manner following ActD treatment (Figure 3B, fifth and fourth row, lane 2-4, respectively and Figure S3).

Accordingly, we shed light on a hindering role of P53 in the ubiquitylation of S2P RNAPII at the early steps of transcription elongation block, therefore protecting it from preliminary degradation.

### 2.4. ATM, DNAPK and P53 coordinate the interaction of WWP2 and CUL3 with S2P RNAPII

It was previously demonstrated that WWP2 interacts both *in vitro* and *in vivo* with the largest subunit (RPB1) of mouse RNAPII regardless of its phosphorylation state and the cellular conditions [32]. This was further supported in human system, in which WWP2 was shown to interact with 11 out of 12 subunits of RNAPII and following DNA damage, the interactions as well as the proteasomal degradation of S2P RNAPII are coordinated by the kinase activity of DNAPK [31]. However, it is important to note that according to the type and the severity of DNA damage, different E3 ligase complexes are involved in the temporal regulation of S2P RNAPII removal and some of them have yet to be unraveled.

These results prompted us to investigate whether the interaction between WWP2 E3 ligase and S2P RNAPII is affected by P53, ATM or DNAPK. Additionally, based on our preliminary unpublished results, we hypothesized a possible, yet unidentified role of CUL3 in the ubiquitylation-mediated removal of S2P RNAPII. To reveal this issue, we immunoprecipitated the S2P RNAPII protein pool in non-treated, 8 h and 24 h ActD treated HCT116 p53+/+ and p53-/- cells following ATMi or DNAPKi treatment and we further investigated whether S2P RNAPII interacted with WWP2 or CUL3 (Figure 4A and Figure 5A, respectively). The immunoprecipitated protein amounts are very diverse among the samples, therefore the density of the detected chemiluminescent signals of either WWP2 or CUL3 was normalised to the adequate signal of S2P RNAPII and then quantified (Figure 4B-C and 5B-C).

**Figure 4.**
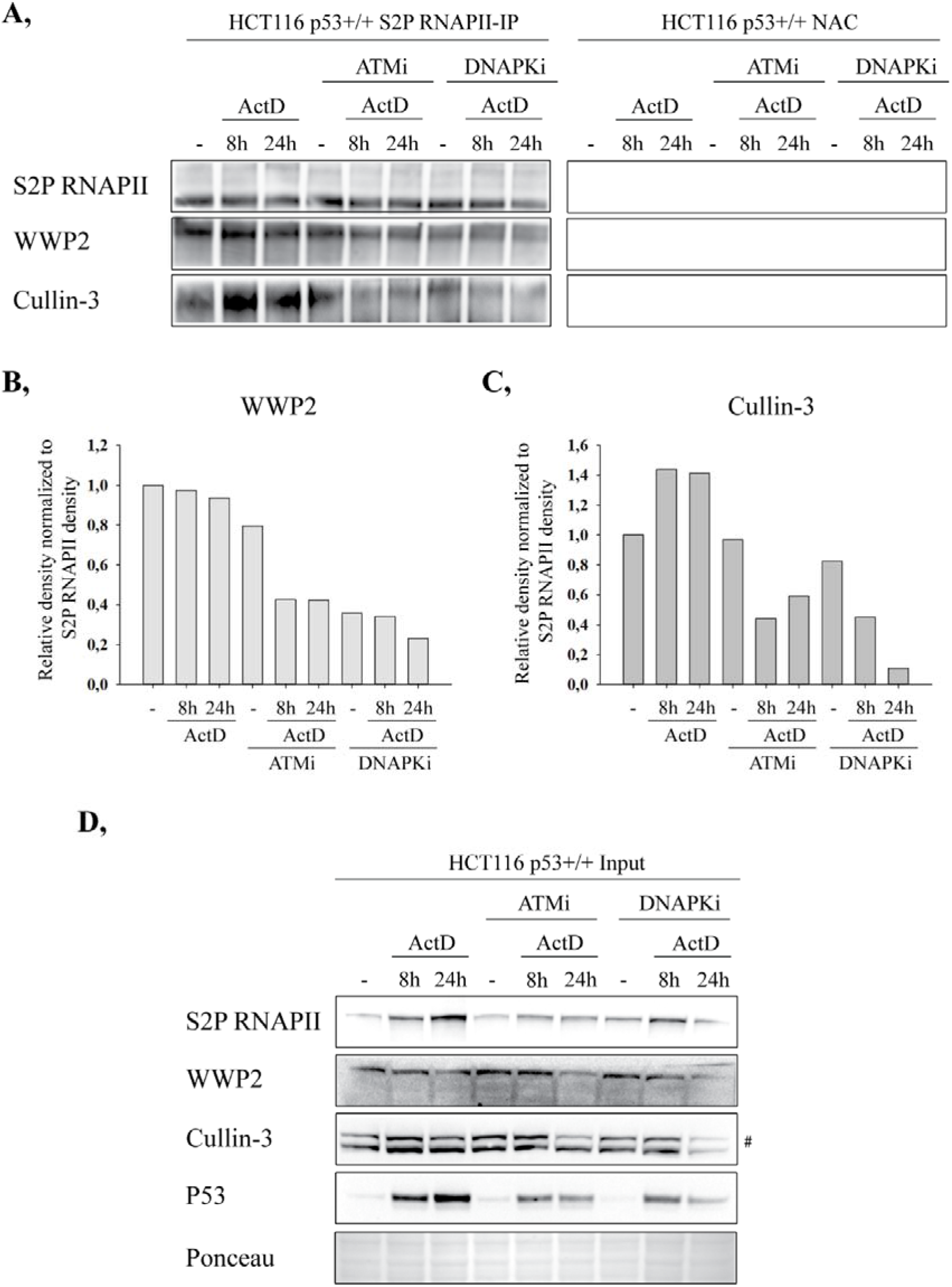
In HCT116 p53+/+ cells, S2P RNAPII interacts with WWP2 and Cullin-3, which is influenced by ATM, DNAPK and P53. **(A)** Co-immunoprecipitation (co-IP) experiment was performed in HCT116 p53+/+ cells. The immunoprecipitation (IP) was accomplished with anti-S2P RNAPII antibody. The efficiency of the IP was demonstrated by anti-S2P RNAPII in Western blot (first row, left panel). Interactions were identified with anti-WWP2 and anti-Cullin-3 antibodies (left panel, second and third row, respectively). Changes in interactions between S2P RNAPII and either WWP2 or Cullin-3 were investigated as a response to 8 h and 24 h ActD treatment in the presence or in the absence of ATMi or DNAPKi (left panel). As IgG control of the IP experiment, no antibody control (NAC) (right panel) from each sample was used, In NACs, no background can be noticed which might be the consequence of the high specificity of the applied anti-S2P RNAPII antibody and Protein A Dynabeads as well. **(B-C)** Density of chemiluminescent signals detected with anti-WWP2 and anti-Cullin-3 antibody in Western blot was normalized to signal densities of the immunoprecipitated S2P RNAPII with Fiji Image J. **(D)** Western blot experiment on HCT116 p53+/+ whole cell lysates (referred to as input samples), which were used for co-IP. Protein level changes of S2P RNAPII, WWP2, Cullin-3 and P53 following 8 h and 24 h ActD treatment in the presence or in the absence of ATMi or DNAPKi was immunodetected using specific antibodies. Ponceau staining was applied to detect the equal loading of the input samples. #: NEDD8-conjugated Cullin-3

**Figure 5.**
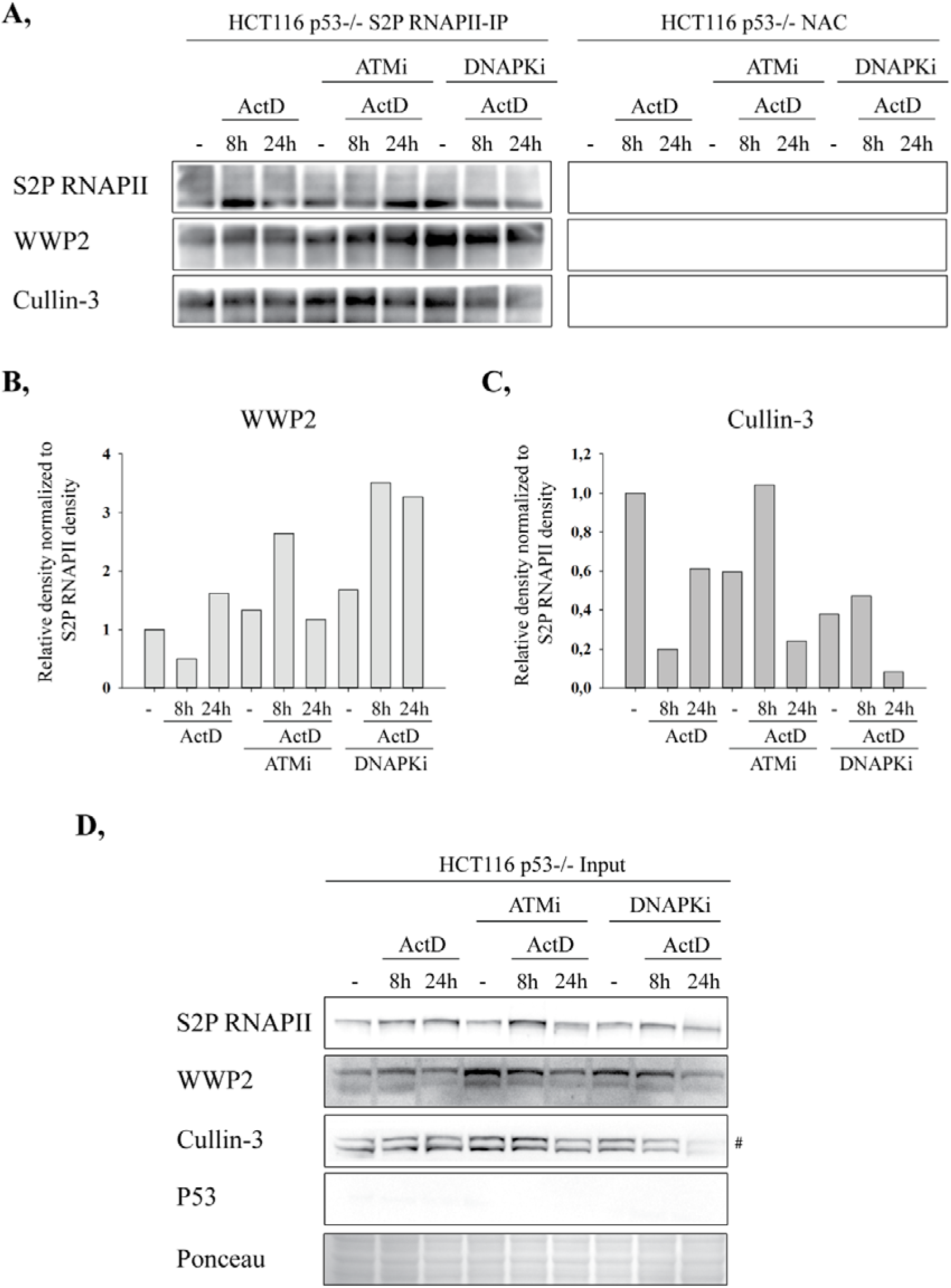
In HCT116 p53-/- cells, S2P RNAPII interacts with WWP2 and Cullin-3, which is mediated by ATM, DNAPK and P53. **(A)** Co-IP experiment was performed in HCT116 p53-/- cells. The immunoprecipitation (IP) was accomplished with anti-S2P RNAPII antibody. The efficiency of the IP was demonstrated by anti-S2P RNAPII in Western blot (left panel, first row). Interactions were identified with anti-WWP2 and anti-Cullin-3 antibodies (left panel, second and third row, respectively). Changes in interactions between S2P RNAPII and either WWP2 or Cullin-3 were investigated as a response to 8 h and 24 h ActD treatment in the presence or in the absence of ATMi or DNAPKi (left panel). As IgG control of the IP experiment, no antibody control (NAC) (right panel) from each sample was used. In NACs, no background can be noticed which might be the consequence of the high specificity of the applied anti-S2P RNAPII antibody and Protein A Dynabeads as well. **(B-C)** Density of chemiluminescent signals detected with anti-WWP2 and anti-Cullin-3 antibody in Western blot was normalized to signal densities of the immunoprecipitated S2P RNAPII with Fiji Image J. **(D)** Western blot experiment on HCT116 p53-/- whole cell lysates (referred to as input samples), which were used for co-IP. Protein level changes of S2P RNAPII, WWP2, Cullin-3 and P53 following 8 h and 24 h ActD treatment in the presence or in the absence of ATMi or DNAPKi was immunodetected using specific antibodies. Ponceau staining was applied to detect the equal loading of the input samples. #: NEDD8-conjugated Cullin-3

In HCT116 p53+/+ cells, interaction between S2P RNAPII and WWP2 is not influenced by stress conditions (Figure 4A, second row, lane 1-3, left panel and Figure 4B, column 1-3), which is consistent with the previous findings published by different laboratories [31,32]. In the lack of ATM activation, the interaction is weakened following ActD treatment (Figure 4A, second row, lane 5-6 compared to lane 4 on left panel and Figure 4B, column 5-6 compared to column 4). Upon DNAPKi, a slight decrease can be observed in S2P RNAPII-WWP2 interaction at 24 h ActD treatment (Figure 4A, second row, lane 9 compared to lane 7 on left panel and Figure 4B, column 9 compared to column 7). Moreover, in all the DNAPKi treated samples we can notice a dramatic decline in the interaction between WWP2 and S2P RNAPII (Figure 4A, second row, lane 7-9 compared to lane 1-3 and Figure 4B, column 7-9 compared to column 1-3). On the other hand, S2P RNAPII and CUL3 interaction is stronger as a response to ActD, while it is greatly reduced following inhibition of either ATM or DNAPK (Figure 4A, third row, left panel and Figure 4C). However, under unstressed conditions, no significant difference can be observed upon either ATM or DNAPK inhibition (Figure 4A, third row, lane 1, 4 and 7, left panel and Figure 4C, column 1, 4 and 7).

In HCT116 p53-/- cells, the interaction between S2P RNAPII and WWP2 or CUL3 is getting weaker following 8 h ActD treatment, then it returns close to the normal level at 24 h ActD (Figure 5A, second and third row, lane 1-3, left panel and Figure 5B-C, column 1-3). Following ATMi treatment, opposite phenomenon can be noticed – a stronger interaction can be detected upon 8 h ActD between S2P RNAPII and WWP2 or CUL3, which is highly reduced at 24 h ActD (Figure 5A, second and third row, lane 4-6, left panel and Figure 5B-C, column 4-6). Nonetheless, DNAPKi differently affects these interactions: (I) S2P RNAPII-WWP2 interaction is getting stronger as a response to 8 h and 24 h ActD (Figure 5A, second row, lane 7-9, left panel and Figure 5B, column 7-9), while (II) S2P RNAPII-CUL3 interaction is dramatically weakened only at 24 h ActD (Figure 5A, third row, lane 9, left panel and Figure 5C, column 9). Furthermore, in the lack of either ATM or DNAPK activity, a reduction can be observed in the interaction between S2P RNAPII and CUL3 under basal conditions (Figure 5A, third row, lane 1, 4 and 7, left panel and Figure 5C, column 1, 4 and 7). As an IgG control, we used no antibody control (NAC) for each sample, in which we did not detect immunoprecipitated proteins and evidently no interaction with WWP2 or CUL3 (Figure 4A, right panel and Figure 5A, right panel).

In the general protein level of S2P RNAPII, upon ActD treatment, a stronger accumulation can be observed in case of HCT116 p53+/+, while a slight increase can be detected in HCT116 p53-/- cells (Figure 4D and 5D, first row, lane 1-3, respectively and Figure S4 and S5). The differences in the S2P RNAPII pattern between the input samples of TUBE (Figure 2B and Figure S2) and co-immunoprecipitation (co-IP) experiment (Figure 4D and 5D, Figure S4 and S5) can be the consequence of different cell harvesting methods. In case of co-IP, non-denaturing conditions were used to avoid the disruption of protein-protein interactions, while in TUBE experiment mechanical cell lysis was applied, which provides a better access to the chromatin-bound fraction. This difference can be also explained by the extremely rapid cell division of HCT116 cells, which implicates an accelerated transcription rate resulting in diverse amounts of S2P RNAPII at the same time-points. Alterations in WWP2 and CUL3 protein levels show the same pattern as detected in Figure 2B (Figure 4D and 5D second and third row, respectively and Figure S4 and S5).

In conclusion, in the presence of P53, the lack of activation of either ATM or DNAPK weakens the interaction between S2P RNAPII and WWP2 or CUL3 (Figure 4A, second and third row, Figure 4B-C, respectively). These results are in concert with the findings revealed by TUBEs pull-down in Figure 2. Moreover, in HCT116 p53-/- cells, following solely 8 h ActD treatment, a very low amount of ub-S2P RNAPII was detected in Figure 2, which is in agreement with the weakened interactions detected between S2P RNAPII and WWP2 or CUL3 (Figure 2A, right panel, lane 2 and Figure S2; Figure 5A, second and third row, left panel, lane 2 and Figure 5B-C, column 2, respectively).

We revealed interactions between S2P RNAPII and WWP2 or CUL3 both under physiological conditions and upon transcription elongation block. We also found that these interactions are strikingly influenced by P53, ATM and DNAPK.

## 3. Discussion

Here, we demonstrate a pivotal role of P53, Ataxia-telangiectasia mutated (ATM) and DNA-dependent protein kinase (DNAPK) in DNA damage-induced transcription silencing. Following transcription arrest, P53 delays the chromatin removal of the stalled elongating RNA polymerase II (S2P RNAPII), supporting its positive effect in transcription elongation. We shed light on a yet to be characterized, emerging role of P53, ATM and DNAPK in the ubiquitylation of S2P RNAPII upon transcription arrest. Following Actinomycin D (ActD)-induced transcription arrest, the activity of ATM and DNAPK is required for maintaining the cellular protein level of WW domain-containing protein 2 (WWP2) and Cullin-3 (CUL3), while P53 counteracts the precocious ubiquitylation of S2P RNAPII. In our study, we demonstrate the interaction between CUL3 and S2P RNAPII, suggesting a possible, yet unexplored function of CUL3 in the ubiquitylation of S2P RNAPII. Moreover, we also reveal that interaction between S2P RNAPII and WWP2 or CUL3 is greatly influenced by the presence of P53, and the activity of ATM and DNAPK.

According to the type of DNA damage and the subsequent fate of RNAPII, various ubiquitylation steps can take place [24]. We previously demonstrated in U2OS cells, that the reduction in S2P RNAPII level following ActD treatment is the consequence of its proteasomal degradation, which requires its pre-ubiquitylation [17]. However, the exact proteins [including E3 ligases, deubiquitylases (DUBs) and scaffold proteins] involved in the resolution of transcription elongation block have yet to be identified in human. E3 ligases involved in the ubiquitylation of RNAPII act synergistically to ensure the most beneficial outcome, which facilitates the recruitment of repair factors to the damage site [2]. Polyubiquitylation is a complex process, in which the linkage type determines the destiny of RNAPII. Neural precursor cell expressed developmentally down-regulated protein 4 (NEDD4) was shown to be one of the key E3 ligases responsible for the monoubiquitylation of RNAPII, and also for its subsequent K63-linked polyubiquitin chain extension [35]. Afterwards, this chain can be trimmed by Ubiquitin carboxyl-terminal hydrolase 2 (Ubp2) (shown in yeast), and eventually processed to K48-linked polyubiquitylation catalyzed by ElonginA/B/C-Cullin-5-RING-box protein 2 (EloA/B/C-CUL5-RBX2) and Von Hippel-Lindau/ElonginB/C-Cullin-2-RING-box protein 1 (VHL/EloB/C-CUL2-RBXl) complexes [24,35,36]. K48-linked chains can also be cropped by Ubiquitin carboxyl-terminal hydrolase 3 (Ubp3) (shown in yeast), hence rescuing RNAPII from subsequent degradation [37]. CUL3, as a scaffold protein, is a core subunit of the BCR (BTB-CUL3-RBX1) E3 ubiquitin ligase complex, which has been shown to be involved in oxidative and electrophilic stress-induced cellular processes so far [38,39]. Here, we assign CUL3 in a novel context of being an interaction partner of S2P RNAPII. This interaction was shown to be stronger following ActD treatment, presuming that CUL3 is a possible factor attending in the ubiquitylation of S2P RNAPII. Moreover, the interaction between CUL3 and S2P RNAPII was greatly diminished upon inhibition of ATM or DNAPK, and also in the absence of P53, supporting the role of ATM, DNAPK and P53 in the ubiquitylation-mediated proteolysis of the stalled S2P RNAPII upon transcription block. This result is in accordance with our data from TUBE pull-down, which revealed a great deduction in the ub-S2P RNAPII level following ActD in the absence of P53 or in the loss of activity of either ATM or DNAPK.

WWP2 E3 ligase has been recently identified as an interaction partner of RNAPII and shown to be involved in the ubiquitylation of RNAPII as a response to DNA damage [31,32]. WWP2-related ubiquitylation-mediated proteasomal degradation of RNAPII is precisely coordinated by the activity of DNAPK [31,33]. However, it still remains elusive how DNAPK can trigger WWP2 and the 26S proteasome to the damage site. A possible explanation of this phenomenon is that a third, still unidentified protein can be involved in this process, which is presumably phosphorylated by DNAPK. In our study, following ActD treatment, a great reduction can be noticed in WWP2 protein level upon inhibition of DNAPK regardless of the P53 presence. Additionally, upon ATM inhibition, a slight reduction in WWP2 level can be also noticed following 24 h ActD treatment. Furthermore, in the absence of P53 or in the lack of either ATM or DNAPK activity, we detected less ubiquitylated S2P RNAPII at 8 h ActD treatment. It presumes an indispensable role of the activity of ATM and DNAPK in the DNA damage-related induction of WWP2, which is an essential component for the ubiquitylation of S2P RNAPII. These data are in concert with previous findings, in which DNAPK was shown to be necessary for the WWP2 and the proteasome recruitment to the damage sites [31]. Nonetheless, in this process, P53 presumably acts differently from ATM and DNAPK. We proved that the limited amount of ub-S2P RNAPII detected in the absence of P53 following 8 h ActD treatment is not the consequence of failure in S2P RNAPII ubiquitylation, but rather a negative regulatory role of P53 in the precocious ubiquitylation-related removal of S2P RNAPII. These data are supported by our chromatin immunoprecipitation (ChIP) results, in which we established that following ActD treatment P53 presence gives rise to a shift in the dwell time of the arrested S2P RNAPII at transcriptionally active coding gene regions. This could be the result of a faster removal of the stalled S2P RNAPII in the absence of P53, leading to the resolution of transcription block at earlier time-points. Based on these, the observed accrual in the general level of S2P RNAPII in the absence of P53 upon 8 h and 24 h ActD treatment might indicate the S2P RNAPII moiety, which is engaged in *de novo* transcription at the break sites necessary for the proper repair. In regards with the above, a potential target of DNAPK which might generate a ‘bridge’ between DNAPK and WWP2 might yet to be identified. Nonetheless, based on our findings this role could be accounted to P53, setting a basis for an intricate hypothetical interplay that needs to be unraveled in the future.

## 4. Materials and Methods

### 4.1. Cell cultures

HCT116 p53+/+ and p53-/- (kindly provided by Prof. Bert Vogelstein, John Hopkins University, Baltimore, MD) isogenic colorectal carcinoma cell lines were used for the experiments [40,41]. Cells were grown in high glucose DMEM (Dulbecco’s Modified Eagle Media; Lonza) supplemented with 8 mM glutamine (Sigma-Aldrich), 1x antibiotic-antimycotic solution (Sigma-Aldrich) and 10% fetal bovine serum (FBS; Lonza). Cells were maintained at 37 °C in humidified atmosphere with 5% CO2.

### 4.2. Treatments

400 nM Actinomycin D (ActD) (Sigma-Aldrich) was used to arrest transcription elongation at different time-points. 10 μM ATMi (KU55933, Sigma-Aldrich) and 20 μM DNAPKi (NU7026, Sigma-Aldrich) were used 1 h prior to ActD treatment to block the activity of ATM and DNAPK, respectively.

### 4.3. Chromatin immunoprecipitation (ChIP)

Cells were fixed with 1% formaldehyde (Sigma-Aldrich) for 10 min, then fixation was halted with 125 mM glycine (Sigma-Aldrich). Cells were centrifuged at 2,000 rpm for 5 min at 4 °C, then they were resuspended in cell lysis buffer (5 mM PIPES pH 8.0, 85 mM KCl, 0.5% NP-40; Sigma-Aldrich) complemented with 1xPIC (Roche) and incubated on ice for 10 min. Pellets were depleted with 5 min centrifugation at 2,000 rpm, 4 °C, resuspended in nuclear lysis buffer (50 mM Tris-HCl pH 8.0, 10 mM EDTA pH 8.0, 0.8% SDS; Sigma-Aldrich) supplemented with 1xPIC (Roche) and incubated on ice for 1 h. Chromatins were sheared 4x 20 sec ON/1 min OFF with Bioruptor Pico sonicator (Diagenode), then diluted four times with dilution buffer (10 mM Tris-HCl pH 8.0, 0.5 mM EGTA pH 8.0, 1% Triton X-100, 140 mM NaCl; Sigma-Aldrich) complemented with 1xPIC (Roche). 30 μg chromatin samples were pre-cleared with 4 μl Sheep anti-Rabbit IgG Dynabeads (Novex) for 2 h rotation at 4 °C. Pre-cleared chromatin samples were incubated with 2 μg anti-S2P RNAPII (Abcam, ab5095) antibody overnight rotating at 4 °C. Chromatin-antibody complexes were captured overnight rotating with 40 μl Sheep anti-Rabbit IgG Dynabeads (Novex). Subsequently to several washing steps, chromatin-antibody complexes were eluted and precipitated. Pellets were resuspended in TE buffer (10 mM Tris-HCl pH 8.0, 1 mM EDTA pH 8.0; Sigma-Aldrich) and reverse-crosslinked. The desired DNA fragments were purified with phenol-chloroform extraction then precipitated with absolute ethanol. Pellets were dissolved in TE buffer. Occupancy of S2P RNAPII was monitored with qPCR (Thermo PikoReal 96 Real-Time PCR system; Thermo Fisher Scientific). Sequences of primers used for qPCR are listed in Table S1.

qPCR quantification was performed by using a TIC (total input control) standard curve. The precipitated amount of DNA in each sample was normalized to the amount of DNA in the NAC (no antibody control).

### 4.4. Western blot

HCT116 p53+/+ and p53-/- cells were harvested in lysis buffer (150 mM NaCl, 1% Triton X-100, 50 mM Tris-HCl pH 8.0; Sigma-Aldrich) supplemented with 1xPIC (Roche), 20 μM DUBi (Calbiochem) and 1x PhosSTOP (Roche) on ice for 10 min, then sonicated 10x 30 sec ON/30 sec OFF in Bioruptor Pico sonicator (Diagenode). Protein concentration was measured with Pierce^™^ BCA Protein Assay Kit (Thermo Fisher Scientific), then 30 μg protein lysates were mixed with NuPAGE^™^ LDS Sample Buffer (4x) (Thermo Fisher Scientific) and boiled for 10 min. Proteins were separated in pre-casted Bolt^™^ 4-12% Bis-Tris Plus gradient gels (Thermo Fisher Scientific), 1x NuPAGE^™^ MOPS SDS Running Buffer (Thermo Fisher Scientific) and 1x NuPAGE^™^ Transfer Buffer (Thermo Fisher Scientific) were used for SDS-PAGE and transfer, respectively. Proteins were transferred onto Amersham Hybond ECL-nitrocellulose membrane (GE Healthcare). Unspecific binding sites of the membranes were blocked with 5% non-fat dry milk-TBST, then the membranes were incubated with primary and horseradish peroxidase (HRP)-conjugated secondary antibodies represented in Table S2 and Table S3. Chemiluminescent detection was conducted using Immobilon Western Chemiluminescent HRP substrate (Millipore) and G:BOX Chemi XRQ (Syngene) system.

### 4.5. Co-immunoprecipitation (co-IP)

HCT116 p53+/+ and p53-/- cells were harvested in non-denaturing lysis buffer (150 mM NaCl, 1% Triton X-100, 50 mM Tris-HCl pH 8.0; Sigma-Aldrich) complemented with 1xPIC (Roche), 20 μM DUBi (Calbiochem) and 1x PhosSTOP (Roche) on ice for 1 h, then protein concentration was measured with Pierce^™^ BCA Protein Assay Kit (Thermo Fisher Scientific). 300 μg lysates were pre-cleared with 2 μl Protein A Dynabeads (Invitrogen) for 1 h at 4 °C, then incubated ON with 2 μg anti-S2P RNAPII (ab5095) antibody. Next day, 10 μl Protein A Dynabeads (Invitrogen) was added to each sample and rotated for 3 h at 4 °C. No antibody controls (NACs) were pre-cleared and incubated with beads only. Beads were washed four times with lysis buffer complemented with 1xPIC (Roche), 20 μM DUBi (Calbiochem) and 1x PhosSTOP (Roche). Protein-protein complexes were eluted in NuPAGE^™^ LDS Sample Buffer (4x) (Thermo Fisher Scientific) at 100 °C for 10 min.

### 4.6. Tandem ubiquitin-binding entities (TUBEs) assay

TUBE2 (UM402; LifeSensors) agarose beads were used to capture polyubiquitin moieties from HCT116 p53+/+ and p53-/- cell lysates. Cells were harvested in TENT buffer (50 mM Tris-HCl pH 8.0, 2 mM EDTA pH 8.0, 150 mM NaCl, 1% Triton X-100; Sigma-Aldrich) complemented with 1xPIC (Roche), 20 μM DUBi (Calbiochem) and 1x PhosSTOP (Roche), incubated on ice for 10 min, then sonicated 12x 30 sec ON/30 sec OFF in Bioruptor Pico sonicator (Diagenode). Afterwards, samples were centrifuged at 14,000g for 10 min at 4 °C. Protein concentration of the supernatants was measured with Pierce^™^ BCA Protein Assay Kit (Thermo Fisher Scientific). 20 μl TUBEs beads/IP were washed twice with TBST (20 mM Tris-HCl pH 8.0, 150 mM NaCl, 0.1% Tween 20; Sigma-Aldrich) complemented with 1xPIC (Roche), 20 μM DUBi (Calbiochem) and 1x PhosSTOP (Roche). Between each washing step, beads were centrifuged at 1,000g for 1 min at 4 °C. 1.5 mg protein was added to 20 μl TUBEs beads and exceeded with TENT + inhibitors up to 500 μl final volume, then samples were rotated for 2 h at 4 °C. Beads-polyubiquitin complexes were washed 3x with TBST + inhibitors, then eluted in 40 μl final volume of the mixture of TENT + inhibitors and NuPAGE^™^ LDS Sample Buffer (4x) (Thermo Fisher Scientific) at 100 °C for 10 min.

### 4.7. Statistics

Statistical analyses were performed to show significant differences between ChIP datasets with one-way ANOVA after checking the normal distribution of the data. Western blots were quantified by using Fiji Image J.

## 5. Conclusions

In this study we reveal an outstanding role of Ataxia-telangiectasia mutated (ATM), DNA-dependent protein kinase (DNAPK) and P53 in the ubiquitylation of the elongating RNA polymerase II (S2P RNAPII) upon transcription elongation block. We found that P53 delays the removal of S2P RNAPII following Actinomycin D (ActD)-induced transcription arrest, confirming the supportive role of P53 in transcription elongation. We showed that ATM and DNAPK have an activator function on WW domain-containing protein 2 (WWP2) E3 ligase involved in the DNA damage-related ubiquitylation of S2P RNAPII, while P53 has a negative impact on the early ubiquitylation of S2P RNAPII. We demonstrated that S2P RNAPII interacts with WWP2 and Cullin-3 (CUL3), a scaffold protein of the BTB-CUL3-RBX1 E3 ligase complex. Furthermore, these interactions are highly mediated by P53 and the kinase activity of ATM and DNAPK. The still unraveled interaction between S2P RNAPII and CUL3 suggests a possible, yet uncharacterized role of CUL3 in the ubiquitylation of S2P RNAPII, which has to be clarified in the future.

## Supplementary Materials

The following are available online at www.mdpi.com/xxx/s1, Figure S1: Changes in S2P RNAPII protein level following ActD treatment at earlier time-points, Figure S2: Relative density of Western blots represented in Figure 2, Figure S3: Relative density of Western blots represented in Figure 3, Figure S4: Relative density of Western blots represented in Figure 4D, Figure S5: Relative density of Western blots represented in Figure 5D, Table S1: Primers used for ChIP-qPCR, Table S2: Antibodies used for protein detection in whole cell lysates by Western blot, and Table S3: Antibodies used for protein detection in immunoprecipitated samples by Western blot.

## Author Contributions

Conceptualization, B.N.B., V.P., T.P.; methodology, B.N.B., V.P., and T.P.; validation, B.N.B., V.P. and T.P.; formal analysis, Z.P. and B.N.B.; investigation, B.N.B., V.P., H.M., Z.U., Z.G.P. and T.P.; resources, I.B., B.N.B. and T.P.; writing—original draft preparation, B.N.B. and T.P.; writing—review and editing, V.P., H.M., Z.U., Z.G.P., I.B. and T.P.; visualization, B.N.B.; supervision, B.N.B. and T.P.; project administration, B.N.B, V.P. and T.P.; funding acquisition, B.N.B and T.P. All authors have read and agreed to the published version of the manuscript.

## Acknowledgments

We are grateful Prof. Dr. Bert Vogelstein for providing HCT116 p53-/- cell line. Additionally, we appreciate Prof. Dr. Evanthia Soutoglou for supporting us with reagents.

## Publisher’s Note

MDPI stays neutral with regard to jurisdictional claims in published maps and institutional affiliations. 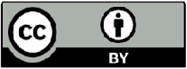 © 2020 by the authors. Submitted for possible open access publication under the terms and conditions of the Creative Commons Attribution (CC BY) license (http://creativecommons.org/licenses/by/4.0/).

## References

1. Beucher, A.; Birraux, J.; Tchouandong, L.; Barton, O.; Shibata, A.; Conrad, S.; Goodarzi, A.A.; Krempler, A.; Jeggo, P.A.; Lobrich, M. ATM and Artemis promote homologous recombination of radiation-induced DNA double-strand breaks in G2. Embo J 2009, 28, 3413–3427, doi:10.1038/emboj.2009.276.

2. Borsos, B.N.; Majoros, H.; Pankotai, T. Ubiquitylation-Mediated Fine-Tuning of DNA Double-Strand Break Repair. Cancers (Basel) 2020, 12, doi:10.3390/cancers12061617.

3. Brandsma, I.; Gent, D.C. Pathway choice in DNA double strand break repair: observations of a balancing act. Genome Integr 2012, 3, 9, doi:10.1186/2041-9414-3-9.

4. Ciccia, A.; Elledge, S.J. The DNA damage response: making it safe to play with knives. Mol Cell 2010, 40, 179–204, doi:10.1016/j.molcel.2010.09.019.

5. Stiff, T.; O’Driscoll, M.; Rief, N.; Iwabuchi, K.; Lobrich, M.; Jeggo, P.A. ATM and DNA-PK function redundantly to phosphorylate H2AX after exposure to ionizing radiation. Cancer Res 2004, 64, 2390–2396, doi:10.1158/0008-5472.can-03-3207.

6. Banin, S.; Moyal, L.; Shieh, S.; Taya, Y.; Anderson, C.W.; Chessa, L.; Smorodinsky, N.I.; Prives, C.; Reiss, Y.; Shiloh, Y., et al. Enhanced phosphorylation of p53 by ATM in response to DNA damage. Science 1998, 281, 1674–1677, doi:10.1126/science.281.5383.1674.

7. Canman, C.E.; Lim, D.S.; Cimprich, K.A.; Taya, Y.; Tamai, K.; Sakaguchi, K.; Appella, E.; Kastan, M.B.; Siliciano, J.D. Activation of the ATM kinase by ionizing radiation and phosphorylation of p53. Science 1998, 281, 1677–1679, doi:10.1126/science.281.5383.1677.

8. Siliciano, J.D.; Canman, C.E.; Taya, Y.; Sakaguchi, K.; Appella, E.; Kastan, M.B. DNA damage induces phosphorylation of the amino terminus of p53. Genes Dev 1997, 11, 3471–3481, doi:10.1101/gad.11.24.3471.

9. Lees-Miller, S.P.; Sakaguchi, K.; Ullrich, S.J.; Appella, E.; Anderson, C.W. Human DNA-activated protein kinase phosphorylates serines 15 and 37 in the amino-terminal transactivation domain of human p53. Mol Cell Biol 1992, 12, 5041–5049, doi:10.1128/mcb.12.11.5041.

10. Finzel, A.; Grybowski, A.; Strasen, J.; Cristiano, E.; Loewer, A. Hyperactivation of ATM upon DNA-PKcs inhibition modulates p53 dynamics and cell fate in response to DNA damage. Mol Biol Cell 2016, 27, 2360–2367, doi:10.1091/mbc.E16-01-0032.

11. Sun, T.; Li, X.; Shen, P. Modeling amplified p53 responses under DNA-PK inhibition in DNA damage response. Oncotarget 2017, 8, 17105–17114, doi:10.18632/oncotarget.15062.

12. Zhou, Y.; Lee, J.H.; Jiang, W.; Crowe, J.L.; Zha, S.; Paull, T.T. Regulation of the DNA Damage Response by DNA-PKcs Inhibitory Phosphorylation of ATM. Mol Cell 2017, 65, 91–104, doi:10.1016/j.molcel.2016.11.004.

13. Chen, B.P.; Uematsu, N.; Kobayashi, J.; Lerenthal, Y.; Krempler, A.; Yajima, H.; Lobrich, M.; Shiloh, Y.; Chen, D.J. Ataxia telangiectasia mutated (ATM) is essential for DNA-PKcs phosphorylations at the Thr-2609 cluster upon DNA double strand break. J Biol Chem 2007, 282, 6582–6587, doi:10.1074/jbc.M611605200.

14. Farmer, G.; Bargonetti, J.; Zhu, H.; Friedman, P.; Prywes, R.; Prives, C. Wild-Type P53 Activates Transcription Invitro. Nature 1992, 358, 83–86, doi:DOI 10.1038/358083a0.

15. Kern, S.E.; Kinzler, K.W.; Bruskin, A.; Jarosz, D.; Friedman, P.; Prives, C.; Vogelstein, B. Identification of P53 as a Sequence-Specific DNA-Binding Protein. Science 1991, 252, 1708–1711, doi:DOI 10.1126/science.2047879.

16. Balakrishnan, S.K.; Gross, D.S. The tumor suppressor p53 associates with gene coding regions and co-traverses with elongating RNA polymerase II in an in vivo model. Oncogene 2008, 27, 2661–2672, doi:10.1038/sj.onc.1210935.

17. Borsos, B.N.; Huliak, L.; Majoros, H.; Ujfaludi, Z.; Gyenis, A.; Pukler, P.; Boros, I.M.; Pankotai, T. Human p53 interacts with the elongating RNAPII complex and is required for the release of actinomycin D induced transcription blockage. Sci Rep 2017, 7, 40960, doi:10.1038/srep40960.

18. Kim, S.; Balakrishnan, S.K.; Gross, D.S. p53 Interacts with RNA polymerase II through its core domain and impairs Pol II processivity in vivo. Plos One 2011, 6, e22183, doi:10.1371/journal.pone.0022183.

19. Woudstra, E.C.; Gilbert, C.; Fellows, J.; Jansen, L.; Brouwer, J.; Erdjument-Bromage, H.; Tempst, P.; Svejstrup, J.Q. A Rad26-Def1 complex coordinates repair and RNA pol II proteolysis in response to DNA damage. Nature 2002, 415, 929–933, doi:DOI 10.1038/415929a.

20. Svejstrup, J.Q. Rescue of arrested RNA polymerase II complexes. J Cell Sci 2003, 116, 447–451, doi:10.1242/jcs.00271.

21. Borsos, B.N.; Majoros, H.; Pankotai, T. Emerging Roles of Post-Translational Modifications in Nucleotide Excision Repair. Cells-Basel 2020, 9, doi:ARTN 1466 10.3390/cells9061466.

22. Charlet-Berguerand, N.; Feuerhahn, S.; Kong, S.E.; Ziserman, H.; Conaway, J.W.; Conaway, R.; Egly, J.M. RNA polymerase II bypass of oxidative DNA damage is regulated by transcription elongation factors. Embo J 2006, 25, 5481–5491, doi:10.1038/sj.emboj.7601403.

23. Golebiowski, F.M.; Li, C.L.; Qnishi, Y.; Samara, N.L.; Sugasawa, K.; Yang, W. Tripartite DNA Lesion Recognition and Verification by XPC, TFIIH, and XPA in Nucleotide Excision Repair. Faseb J 2016, 30.

24. Wilson, M.D.; Harreman, M.; Svejstrup, J.Q. Ubiquitylation and degradation of elongating RNA polymerase II: The last resort. Bba-Gene Regul Mech 2013, 1829, 151–157, doi:10.1016/j.bbagrm.2012.08.002.

25. Mitsui, A.; Sharp, P.A. Ubiquitination of RNA polymerase II large subunit signaled by phosphorylation of carboxyl-terminal domain. P Natl Acad Sci USA 1999, 96, 6054–6059, doi:DOI 10.1073/pnas.96.11.6054.

26. Rockx, D. A.P.; Mason, R.; van Hoffen, A.; Barton, M.C.; Citterio, E.; Bregman, D.B.; van Zeeland, A.A.; Vrieling, H.; Mullenders, L.H.F. UV-induced inhibition of transcription involves repression of transcription initiation and phosphorylation of RNA polymerase II. P Natl Acad Sci USA 2000, 97, 10503–10508, doi:DOI 10.1073/pnas.180169797.

27. Anindya, R.; Aygun, O.; Svejstrup, J.Q. Damage-induced ubiquitylation of human RNA polymerase II by the ubiquitin ligase Nedd4, but not Cockayne syndrome proteins or BRCA1. Molecular Cell 2007, 28, 386–397, doi:10.1016/j.molcel.2007.10.008.

28. Kuznetsova, A.V.; Meller, J.; Schnell, P.O.; Nash, J.A.; Ignacak, M.L.; Sanchez, Y.; Conaway, J.W.; Conaway, R.C.; Czyzyk-Krzeska, M.F. von Hippel-Lindau protein binds hyperphosphorylated large subunit of RNA polymerase II through a proline hydroxylation motif and targets it for ubiquitination. P Natl Acad Sci USA 2003, 100, 2706–2711, doi:10.1073/pnas.0436037100.

29. Yasukawa, T.; Kamura, T.; Kitajima, S.; Conaway, R.C.; Conaway, J.W.; Aso, T. Mammalian Elongin A complex mediates DNA-damage-induced ubiquitylation and degradation of Rpb1. Embo J 2008, 27, 3256–3266, doi:10.1038/emboj.2008.249.

30. Starita, L.M.; Horwitz, A.A.; Keogh, M.C.; Ishioka, C.; Parvin, J.D.; Chiba, N. BRCA1/BARD1 ubiquitinate phosphorylated RNA polymerase II. J Biol Chem 2005, 280, 24498–24505, doi:10.1074/jbc.M414020200.

31. Caron, P.; Pankotai, T.; Wiegant, W.W.; Tollenaere, M.A.X.; Furst, A.; Bonhomme, C.; Helfricht, A.; de Groot, A.; Pastink, A.; Vertegaal, A.C.O., et al. WWP2 ubiquitylates RNA polymerase II for DNA-PK-dependent transcription arrest and repair at DNA breaks. Gene Dev 2019, 33, 684–704, doi:10.1101/gad.321943.118.

32. Li, H.; Zhang, Z.H.; Wang, B.B.; Zhang, J.M.; Zhao, Y.M.; Jin, Y. Wwp2-mediated ubiquitination of the RNA polymerase II large subunit in mouse embryonic pluripotent stem cells. Mol Cell Biol 2007, 27, 5296–5305, doi:10.1128/Mcb.01667-06.

33. Pankotai, T.; Bonhomme, C.; Chen, D.; Soutoglou, E. DNAPKcs-dependent arrest of RNA polymerase II transcription in the presence of DNA breaks. Nat Struct Mol Biol 2012, 19, 276–U229, doi:10.1038/nsmb.2224.

34. Wimuttisuk, W.; Singer, J.D. The Cullin3 ubiquitin ligase functions as a Nedd8-bound heterodimer. Mol Biol Cell 2007, 18, 899–909, doi:10.1091/mbc.e06-06-0542.

35. Harreman, M.; Taschner, M.; Sigurdsson, S.; Anindya, R.; Reid, J.; Somesh, B.; Kong, S.E.; Banks, C.A.S.; Conaway, R.C.; Conaway, J.W., et al. Distinct ubiquitin ligases act sequentially for RNA polymerase II polyubiquitylation. P Natl Acad Sci USA 2009, 106, 20705–20710, doi:10.1073/pnas.0907052106.

36. Kee, Y.; Lyon, N.; Huibregtse, J.M. The Rsp5 ubiquitin ligase is coupled to and antagonized by the Ubp2 deubiquitinating enzyme. Embo J 2005, 24, 2414–2424, doi:10.1038/sj.emboj.7600710.

37. Kvint, K.; Uhler, J.P.; Taschner, M.J.; Sigurdsson, S.; Erdjument-Bromage, H.; Tempst, P.; Svejstrup, J.Q. Reversal of RNA polymerase II ubiquitylation by the ubiquitin protease ubp3. Molecular Cell 2008, 30, 498–506, doi:10.1016/j.molcel.2008.04.018.

38. Dubiel, W.; Dubiel, D.; Wolf, D.A.; Naumann, M. Cullin 3-Based Ubiquitin Ligases as Master Regulators of Mammalian Cell Differentiation. Trends Biochem Sci 2018, 43, 95–107, doi:10.1016/j.tibs.2017.11.010.

39. Taguchi, K.; Motohashi, H.; Yamamoto, M. Molecular mechanisms of the Keap1-Nrf2 pathway in stress response and cancer evolution. Genes to Cells 2011, 16, 123–140, doi:10.1111/j.1365-2443.2010.01473.x.

40. Bunz, F.; Dutriaux, A.; Lengauer, C.; Waldman, T.; Zhou, S.; Brown, J.P.; Sedivy, J.M.; Kinzler, KW.; Vogelstein, B. Requirement for p53 and p21 to sustain G(2) arrest after DNA damage. Science 1998, 282, 1497–1501, doi:DOI 10.1126/science.282.5393.1497.

41. Sur, S.; Pagliarini, R.; Bunz, F.; Rago, C.; Diaz, L.A.; Kinzler, K.W.; Vogelstein, B.; Papadopoulos, N. A panel of isogenic human cancer cells suggests a therapeutic approach for cancers with inactivated p53. P Natl Acad Sci USA 2009, 106, 3964–3969, doi:10.1073/pnas.0813333106.

