## Supplementaries for "Old players in new posts: the role of P53, ATM and DNAPK in DNA damage-related ubiquitylation-dependent removal of S2P RNAPII"

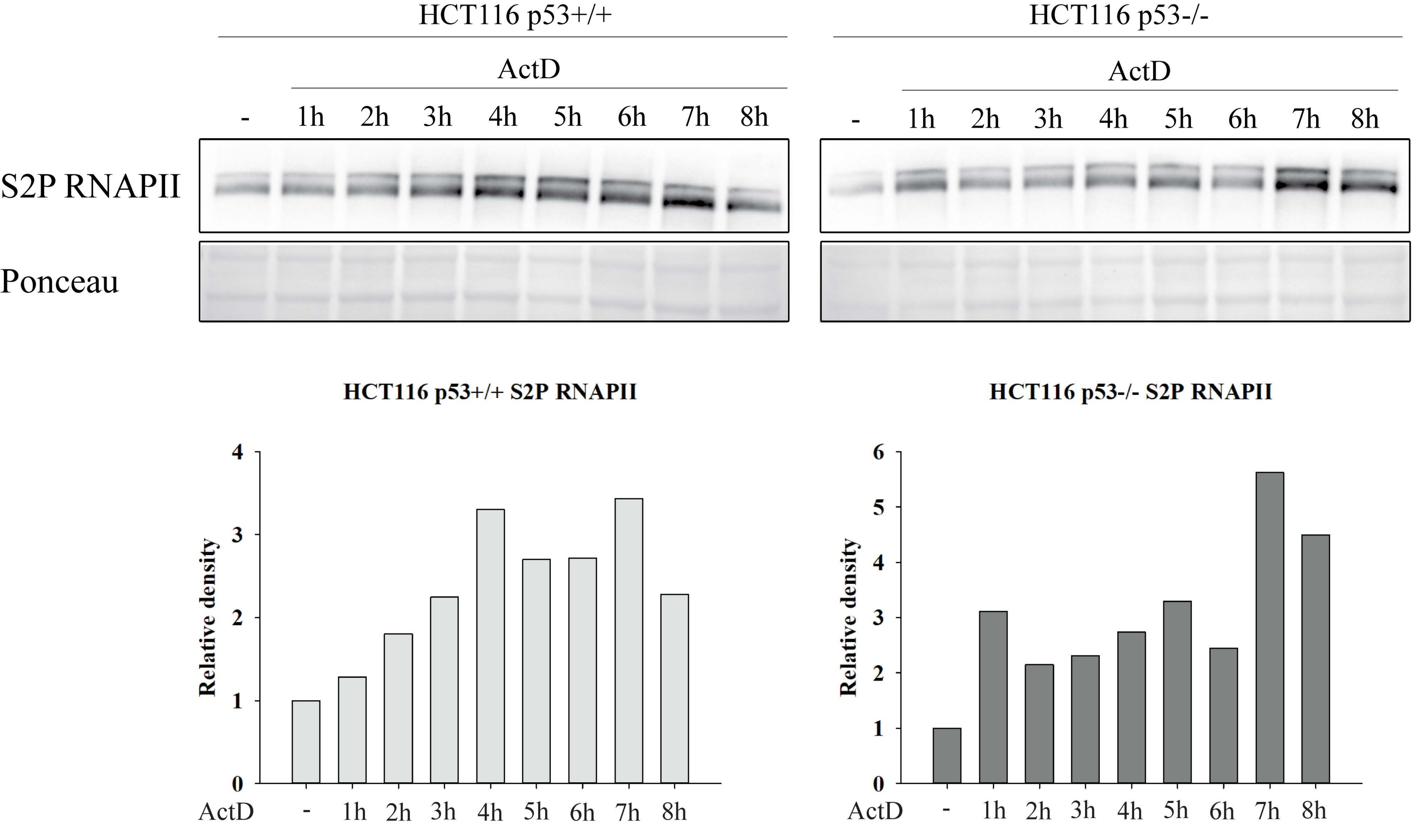


**Figure S1.** Changes in S2P RNAPII protein level following ActD treatment at earlier time-points. Western blot detection of S2P RNAPII protein level upon 1-8 h ActD treatment. Equal loading of the samples was controlled by Ponceau staining. Relative density is shown below the corresponding Western blot.

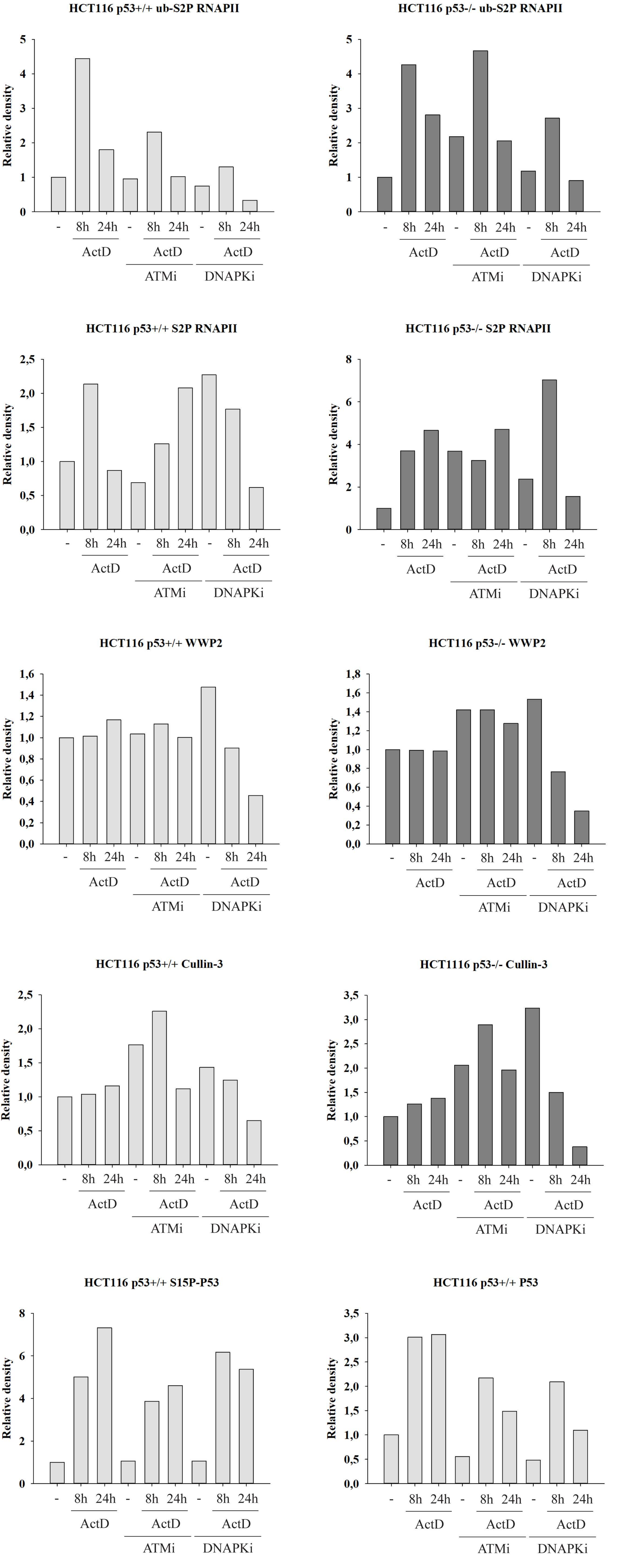


**Figure S2.** Relative density of Western blots represented in Figure 2

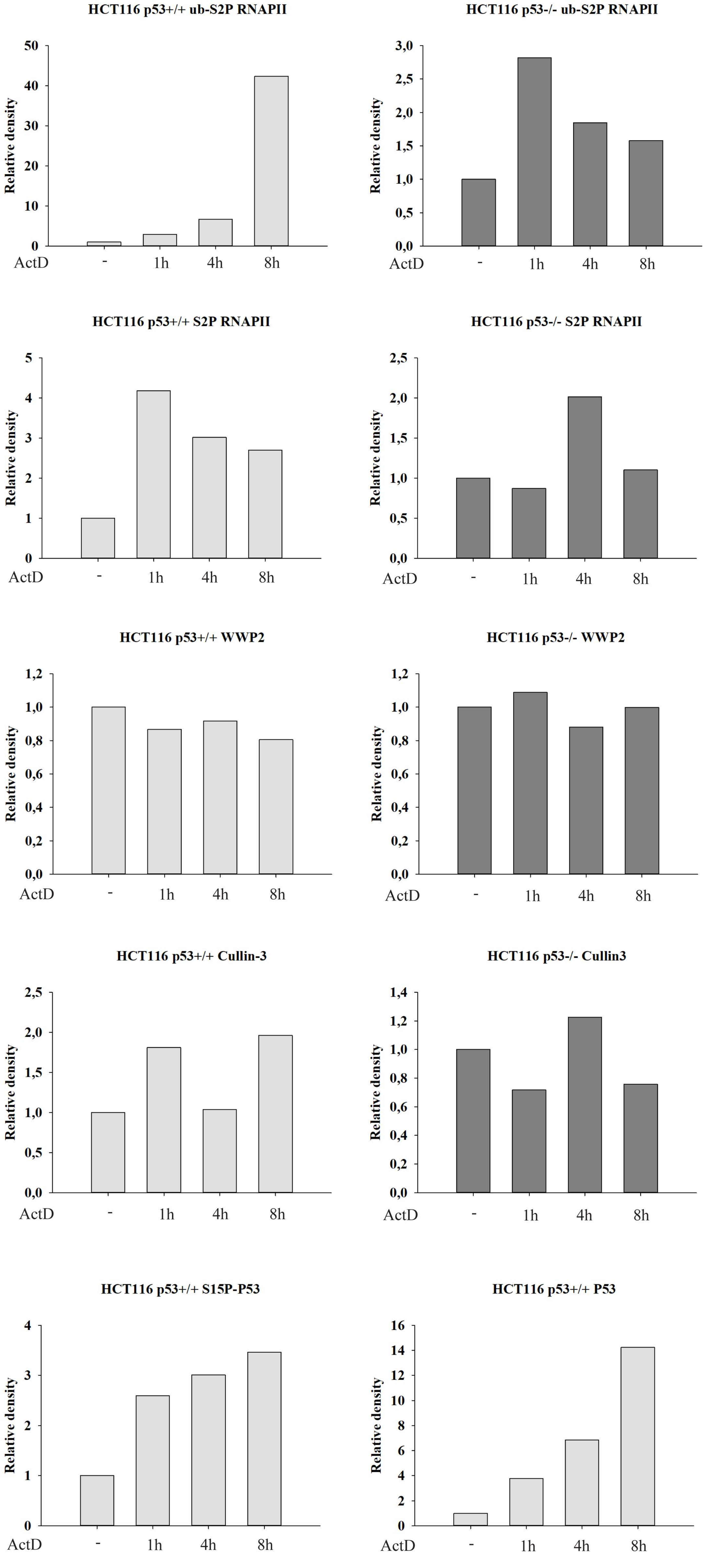


**Figure S3.** Relative density of Western blots represented in Figure 3

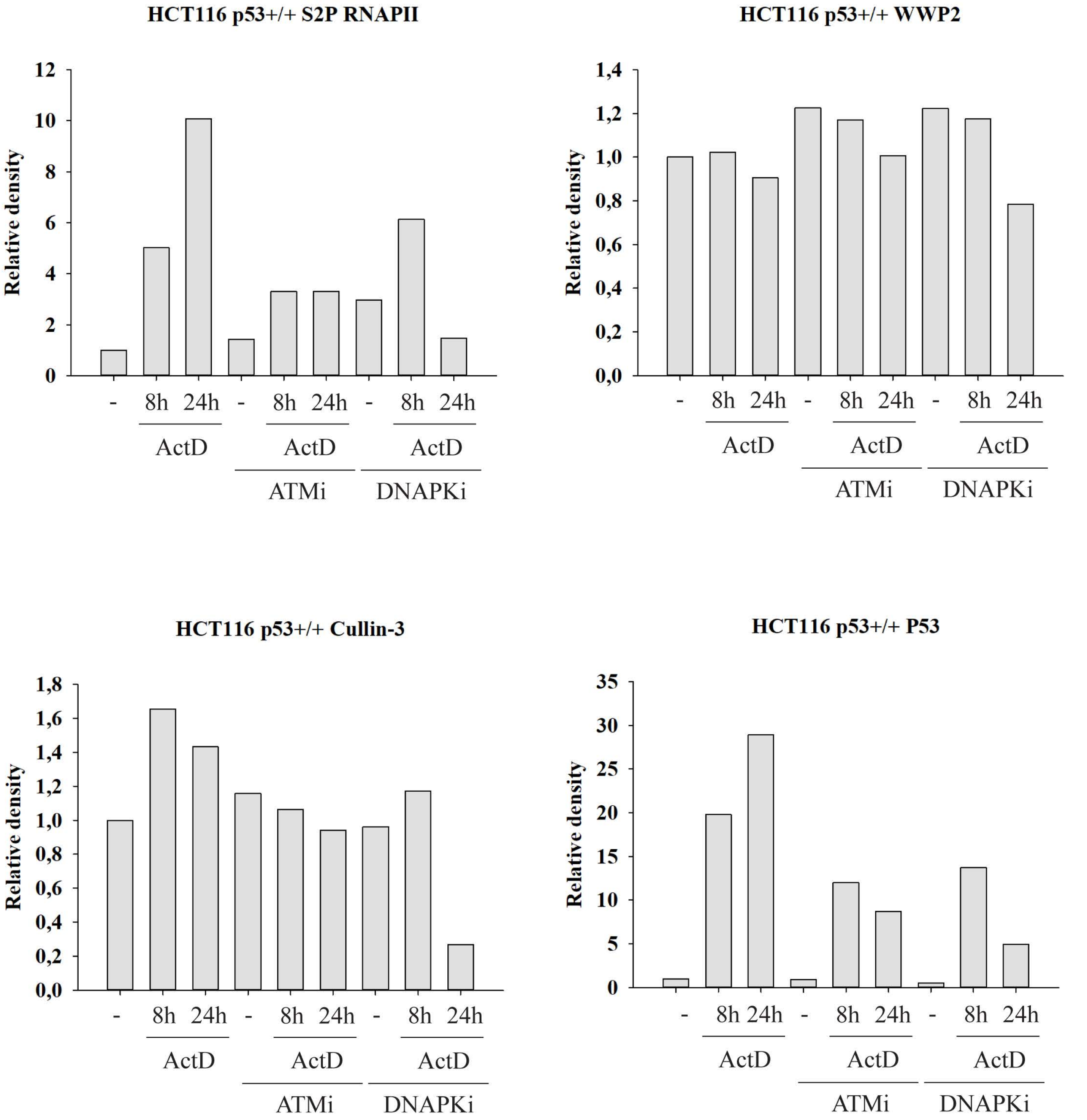


**Figure S4.** Relative density of Western blots represented in Figure 4D

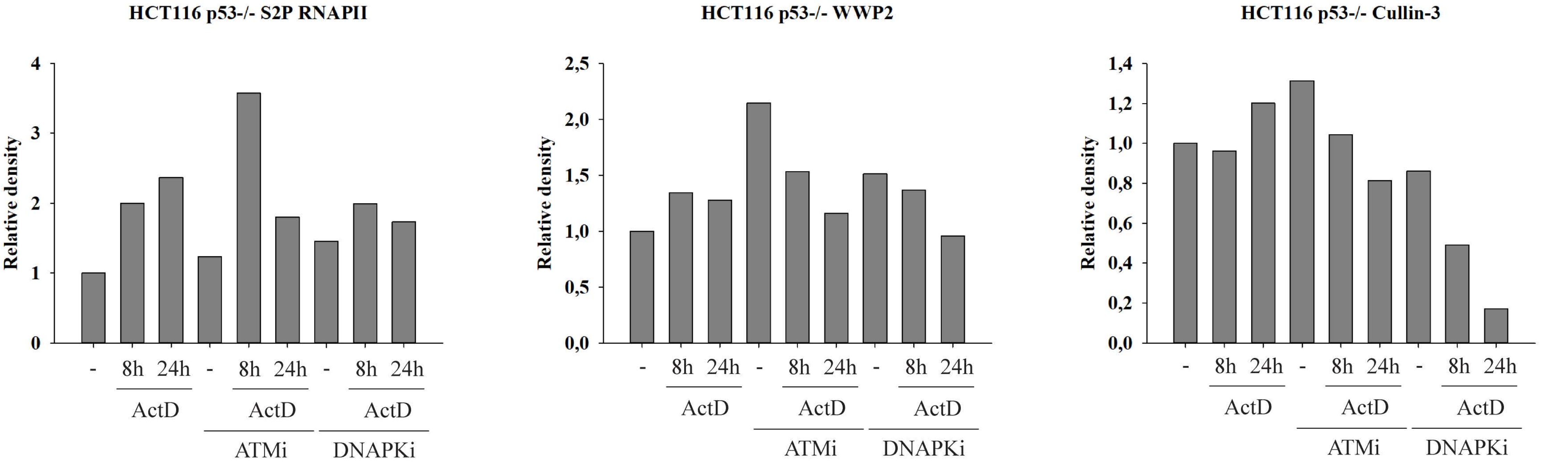


**Figure S5.** Relative density of Western blots represented in Figure 5D

| Gene region | Forward primer (5'-3') | Reverse primer (5'-3') |
|---|---|---|
| *ActB Exon1* | GGCATGGGTCAGAAGGATT | AGGTGTGGTGCCAGATTTTC |
| *ActB Exon2* | TCCCTGGAGAAGAGCTACGA | AGGAAGGAAGGCTGGAAGAG |
| *ActB Intron* | ATGAGGCTGGTGTAAAGCGG | TAAGTGTGCTGGGGTCTTGG |
| *Cdk12 Exon1* | TCTTCCACAGCAACCACCTC | TGAGTGAGTAGAAGGGGGCA |
| *Cdk12 Exon2* | CTTTGTGGTAGCCCTTGTCC | GACGCCTTCGATATTGCTTC |
| *Cdk12 Intron* | CCCCAGGTGAGCTATTTGTC | CAACTGAAGACCCCACCACT |
| *Intergenic region* | TGGAACTTCTGGAAGACACTG | TACACCACTCAAGGGAAACTG |

**Table S1.** Primers used for ChIP-qPCR

| First antibody | | Secondary antibody | |
|---|---|---|---|
| **Type** | **Dilution** | **Type** | **Dilution** |
| anti-S2P RNAPII ab5095 (Abcam) | 1:4000 | GAR-HRP IgG P0448 (DAKO) | 1:16000 |
| anti-CUL3 ab194584 (Abcam) | 1:1000 | GAR-HRP IgG P0448 (DAKO) | 1:2000 |
| anti-WWP2 A302-935A (Bethyl Lab) | 1:1000 | GAR-HRP IgG P0448 (DAKO) | 1:2000 |
| anti-P53 MA5-12557 (Thermo Fisher Scientific) | 1:1000 | RAM-HRP IgG P0260 (DAKO) | 1:2000 |
| anti-S15P P53 9284L (Cell Signaling) | 1:1000 | GAR-HRP IgG P0448 (DAKO) | 1:4000 |

**Table S2.** Antibodies used for protein detection in whole cell lysates by Western blot

| First antibody | | Secondary antibody | |
|---|---|---|---|
| **Type** | **Dilution** | **Type** | **Dilution** |
| anti-S2P RNAPII ab5095 (Abcam) | 1:2000 | GAR-HRP IgG P0448 (DAKO) | 1:8000 |
| anti-CUL3 ab194584 (Abcam) | 1:500 | GAR-HRP IgG P0448 (DAKO) | 1:1000 |
| anti-WWP2 A302-935A (Bethyl Lab) | 1:1000 | GAR-HRP IgG P0448 (DAKO) | 1:4000 |

**Table S3.** Antibodies used for protein detection in immunoprecipitated samples by Western blot
